# Contributions of P_D1_ and P_D2_ to the [P_D1_P_D2_]^+^*-minus-*[P_D1_P_D2_] difference spectrum in the Soret region in Photosystem II

**DOI:** 10.1101/2024.02.15.580457

**Authors:** Alain Boussac, Miwa Sugiura, Ryo Nagao, Takumi Noguchi, A. William Rutherford, Julien Sellés

## Abstract

Flash-induced absorption changes in the Soret region, which originate from the [P_D1_P_D2_]^+^ state, the chlorophyll cation radical formed upon Photosystem II (PSII) excitation, were investigated in Mn-depleted Photosystem II. In wild-type PSII from *Thermosynechococcus elongatus*, the [P_D1_P_D2_]^+^-*minus*-[P_D1_P_D2_] difference spectrum shows a main negative feature at 434 nm and a smaller negative feature at 446 nm [Boussac et al. Photosynth Res (2023), https://doi.org/10.1007/s11120-023-01049-3]. While the main feature at 434 nm is associated with P_D1_^+^ formation, the origin of the dip at 446 nm remains to be identified. For that, we have compared the [P_D1_P_D2_]^+^-*minus*-[P_D1_P_D2_] difference spectra from the PsbA3/H198Q PSII mutant in *T. elongatus* and D2/H197A PSII mutant in *Synechocystis* sp. PCC 6803 with their respective wild type strains. By modifying the P_D1_ axial ligand with the H198Q mutation in the D1 protein in *T. elongatus*, the contribution at 434 nm was shifted to 431 nm, while the contribution at 446 nm was hardly affected. In *Synechocystis* sp. PCC 6803, by modifying the P_D2_ axial ligand with the H197A mutation in the D2 protein, the contribution at 446 nm was downshifted by ∼ 3 nm to ∼ 443 nm, while the main contribution at 432 nm was only slightly shifted upwards to 433 nm. This result suggests that the bleaching seen at 446 nm involves P_D2_. This could reflects a change in the [P_D1_^+^P_D2_]⟷[P_D1_P_D2_^+^] equilibrium or a more complex mechanism.

## Introduction

Oxygenic photosynthesis is responsible for most of the energy input to life on Earth. This process converts the solar energy into fiber, food and fuel, and occurs in cyanobacteria, algae and plants. Photosystem II (PSII), the water-splitting enzyme, is at the heart of this process, see (Shevela et al. 2023) for a recent review.

Mature cyanobacterial PSII generally consists of 20 subunits with 17 trans-membrane and 3 extrinsic membrane proteins. PSII binds 35 chlorophylls *a* (Chl-*a*), 2 pheophytins (Phe), 1 membrane b-type cytochrome, 1 extrinsic c-type cytochrome, 1 non-heme iron, 2 plastoquinones (Q_A_ and Q_B_), the Mn_4_CaO_5_ cluster, 2 Cl^−^, 12 carotenoids and 25 lipids (Suga et al. 2015). The 4^th^ extrinsic PsbQ subunit was also found in PSII from *Synechocystis* sp. PCC 6803 in addition to PsbV, PsbO and PsbU (Gisriel et al. 2022). Most of the Chl bound to PSII have histidine ligands to their Mg^2+^ ions, including P_D1_ and P_D2_ with the D1/H198 and D2/H197, respectively (Suga et al. 2015).

Solar energy conversion into chemical energy starts with the absorption of a photon by a Chl in the antenna forming an excited state. The excitation energy is then transferred to other chlorophylls until it reaches the key pigments in photochemical reaction centre, *e.g*. (Mirkovic et al. 2017) for a review, with 4 Chl-*a* molecules, P_D1_, P_D2_, Chl_D1_, Chl_D2_ and 2 Phe-*a* molecules, Phe_D1_ and Phe_D2_, *e.g*. (Cardona et al. 2012, Holzwarth et al. 2006, Sirohiwal and Pantazis 2023, Romero et al. 2017, Yoneda et al. 2022).

A few picoseconds after the excitation reaches Chl_D1_, a charge separation occurs resulting in the formation of the Chl_D1_^+^Phe_D1_^−^ and then of the [P_D1_P_D2_]^+^Phe_D1_^−^ radical pair states, with the positive charge mainly located on P_D1_, *e.g*. (Holzwarth et al. 2006, Romero et al. 2017, Sirohiwal and Pantazis 2023). After the formation of [P_D1_P_D2_]^+^Phe_D1_^−^, the electron on Phe_D1_^−^ is transferred to Q_A_, the primary quinone electron acceptor, and then to Q_B_, the second quinone electron acceptor. While Q_A_ is only singly reduced under normal conditions, Q_B_ accepts two electrons and two protons before leaving its binding site and being replaced by an oxidized plastoquinone molecule from the membrane plastoquinone pool, *e.g*. (de Causmaecker et al. 2019). On the donor side of PSII, P_D1_^+^ oxidizes Tyr_Z_, the Tyr161 of the D1 polypeptide. The Tyr_Z_^•^ radical is then reduced by the Mn_4_ CaO_5_ cluster, *e.g*. (Shevela et al. 2023) for a recent review. After four charge separations, the Mn_4_CaO_5_ cluster accumulates four oxidizing equivalents and thus cycles through five redox states denoted S_0_ to S_4_. Upon formation of the S_4_-state, two molecules of water are oxidized, the S_0_-state is regenerated and O_2_ is released (Joliot et al. 1969, Kok et al. 1970).

The [P_D1_P_D2_]^+^-*minus*-[P_D1_P_D2_] difference spectrum is much less studied and understood in the Soret region than in the Q_x_ and Q_y_ regions, *e.g*. (Krausz 2013, Reimers et al. 2013). For example, in wild-type PSII, when Tyr_D_^•^ is present, an additional signal in the [PD_1_ P D_2_]^+^-*minus*-[P_D1_P_D2_] difference spectrum was recently observed when compared to the first flash when Tyr_D_ is not oxidized (Boussac et al. 2023). The additional feature was “W-shaped” with troughs at 434 nm and 446 nm. This double trough feature in the [P_D1_P_D2_]^+^-*minus*-[P_D1_P_D2_] difference spectrum was seemingly unmodified in all of the mutants studied except in the P_D2_ mutant, D2/H197A, in *Synechoscystis* sp. PCC 6803 in which the trough at 446 was downshifted by ∼ 3 nm. In this study, however, we lacked the [P_D1_P_D2_]^+^-*minus*-[P_D1_P_D2_] difference spectrum in wild-type *Synechoscystis sp* PCC 6803, which would have been the optimum spectrum for the comparive study. We have therefore measured this difference spectrum in the present work. The [P_D1_P_D2_]^+^-*minus*-[P_D1_P_D2_] difference spectrum in the PsbA3/H198Q mutant was only reported in our previous work in O_2_-evolving PSII at low pH values (Boussac et al. 2023). Therefore, we have also measured this spectrum in Mn-depleted PsbA3/H198Q-PSII at pH 8.6, as for the other spectra, for a better comparison although the Mn-depletion seemed to have no detectable effect on these spectra.

## Materials and Methods

### PSII samples

The His-tagged PSII samples from *T. elongatus* used in this study, PsbA1-PSII, PsbA3-PSII and PsbA3/H198Q-PSII, were purified as described previously (Boussac et al. 2023). The His-tagged PSII samples from *Synechocystis sp* PCC 6803 used in this study, D2/H197A-PSII and WT-PSII (Hayase et al. 2023), were purified as described previously (Boussac et al. 2023). The Mn-depletion procedure has also been described previously (Boussac et al. 2023). In the last step, all the PSII samples were suspended in 1 M betaine, 15 mM CaCl_2_, 15 mM MgCl_2_, 100 mM Tris, pH 8.6.

### UV-visible time-resolved absorption change spectroscopy

Absorption change measurements were performed with a modified lab-built spectrophotometer (Béal et al. 1999) described in detail in (Boussac et al. 2023).

For the ΔI/I measurements, the Mn-depleted PSII samples were diluted in a medium with 1 M betaine, 15 mM CaCl_2_, 15 mM MgCl_2_, and 100 mM Tris with the pH adjusted with HCl at pH 8.6. All the PSII samples were dark-adapted for ∼ 3-4 h at room temperature (20– 22°C) before the addition of 0.1 mM phenyl *p*–benzoquinone (PPBQ) dissolved in dimethyl sulfoxide. In all cases, the chlorophyll concentration of the samples was ∼ 25 µg of Chl mL^−1^. After the ΔI/I measurements, the absorption of each diluted batch of samples was precisely controlled to avoid errors due to the dilution of concentrated samples and the ΔI/I values shown in the figures were normalized to *A*_673_ = 1.75, with ε ∼ 70 mM^−1^·cm^−1^ at 674 nm for dimeric *T. elongatus* PSII (Müh and Zouni 2005).

## Results and Discussion

The two Panels in Fig. 1 show the averaged [P_D1_P_D2_]^+^-*minus*-[P_D1_P_D2_] difference spectra measured 20 ns after the 5^th^ to 10^th^ flashes flash illumination. The 4 first flashes were given to fully oxidize Tyr_D_ (Boussac et al. 2023).

**Figure 1:**
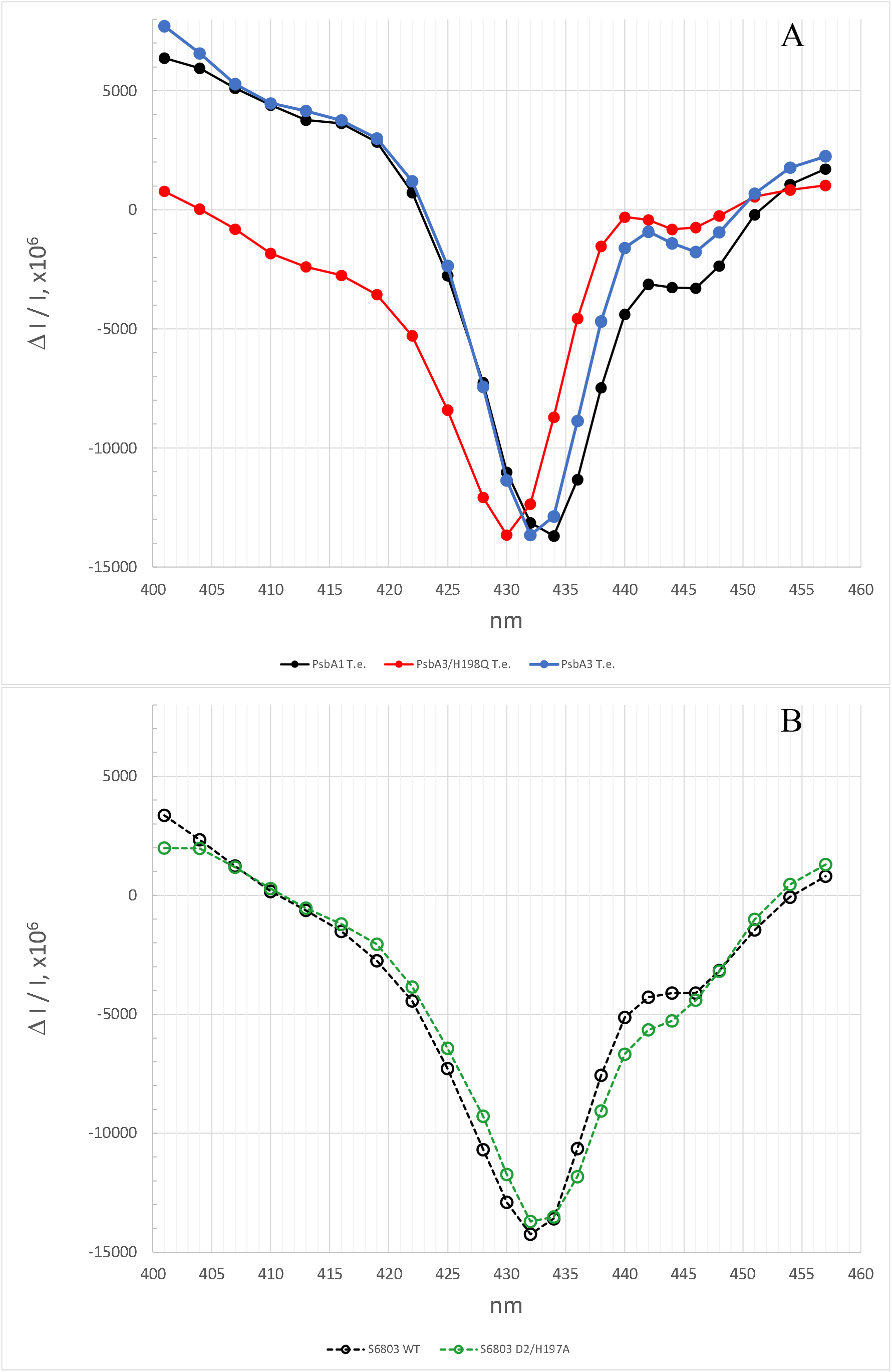
Panel A, averaged spectra recorded 20 ns after the 5^th^ to 10^th^ laser flash illumination at pH 8.6 in Mn-depleted PsbA1-PSII (black spectrum), Mn-depleted PsbA3-PSII (blue spectrum), and PsbA3/H198Q-PSII from *T. elongatus*. Panel B, averaged spectra recorded 20 ns after the 5^th^ to 10^th^ laser flash illumination at pH 8.6 in Mn-depleted PSII (black spectrum), and Mn-depleted D2/H198Q PSII from *Synechocystis* 6803. The Chl concentration was 25 µg mL^−1^ and 100 µM PPBQ was added before the measurements.

Panel A in Fig. 1 shows the spectra in PsbA1-PSII (black spectrum with closed symbols), PsbA3-PSII (blue spectrum with closed symbols), and PsbA3/H198Q-PSII (red spectrum with closed symbols). Two observations, already reported (Boussac et al. 2023), can be made: *i*) the nature of PsbA, *i.e*. PsbA1 *vs* PsbA3, hardly affects the [P_D1_P_D2_]^+^-*minus*-[P_D1_P_D2_] difference spectrum, and *ii*) the largest trough in the [P_D1_P_D2_]^+^-*minus*-[P_D1_P_D2_] difference spectrum in the PsbA3/H198Q-PSII mutant is shifted towards the blue by ∼ 3 nm, as already reported both in *Synechocystis* 6803 and *T. elongatus* (Diner et al. 2001, Sugiura et al. 2016), whereas the minor bleaching at 446 nm is almost unaffected.

Panel B in Fig. 1 shows the [P_D1_P_D2_]^+^-*minus*-[P_D1_P_D2_] difference spectra in the wild-type *Synechocystis* 6803 PSII (black spectrum with open symbols) and D2/H197A mutant PSII (green spectrum with open symbols). Again, two main observations can be made: *i*) the largest trough in the [P_D1_P_D2_]^+^-*minus*-[P_D1_P_D2_] difference spectrum in the wild-type PSII is at ∼ 432 nm while in the D2/H197A mutant is slightly shifted to the red by ∼ 1 nm to ∼ 433 nm, and *ii*) the trough at ∼ 446 nm in the wild-type PSII is more significantly shifted to the blue at ∼ 443 nm in the mutant.

Diner et al. (2001) have reported that the [P_D1_P_D2_]^+^-*minus*-[P_D1_P_D2_] difference spectrum was red shifted by ∼ 1 nm from ∼ 432 nm to ∼ 433 nm due to the D2/H197A mutation. Fig. 2 shows on the same graph the [P_D1_P_D2_]^+^-*minus*-[P_D1_P_D2_] difference spectra in wild-type *Synechocystis* 6803 (black spectrum with open symbols) and the PsbA1-PSII from *T. elongatus* (black spectrum with closed symbols). This comparison clearly shows that the main trough of the spectrum in wild-type *Synechocystis* 6803 is slightly blue-shifted when compared to the spectrum in *T. elongatus* (in contrast to the trough at 446 nm that is not affected). As we did not used the proper control spectrum in (Boussac et al. 2023), *i.e*. the wild-type PSII from *Synechocystis* 6803, the small red shift due to the D2/H197A mutation escaped our detection.

**Figure 2:**
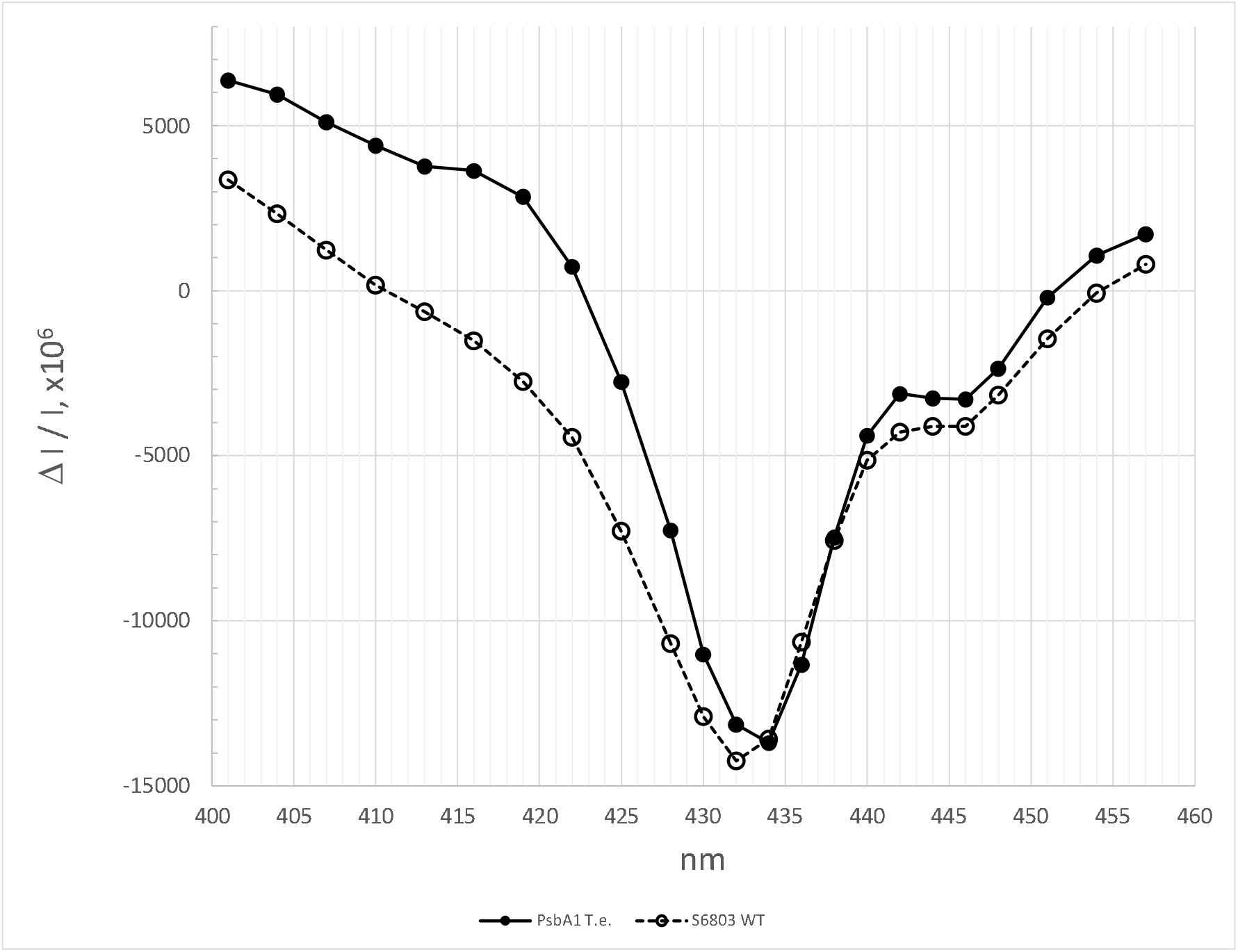
Replot of the spectrum in PsbA1-PSII from *T. elongatus* (closed symbols, full line) and in the wild-type PSII from *Synechocystis* 6803 (open symbols, dashed line).

The P_D2_ mutant, D2/H197A, is the only one in which the trough at 446 nm in the [P_D1_P_D2_]^+^-*minus*-[P_D1_P_D2_] difference spectrum is significantly affected. This supports the hypothesis that the bleaching seen at 446 nm involves P_D2_. The two spectral features could reflects the [P_D1_^+^P_D2_]⟷[P_D1_P_D2_^+^] equilibrium. The relative amplitudes of the trough at 446 nm and 434 nm are indeed in agreement with the relative proportions of PD_1_ ^+^ and PD_2_^+^ in the [P_D1_P_D2_]^+^ cation, roughly 80% and 20%, respectively, from experimental estimates, *e.g*. (Rigby et al. 1994, Diner et al. 2001, Okubo et al. 2007, Nagao et al. 2017). Alternatively, it could be due to something more complex due to couplings between several pigments or charge fluctuations between them (Narzi et. 2016). These data could be useful for computational works aiming at understanding the [P_D1_P_D2_]^+^-*minus*-[P_D1_P_D2_] difference spectrum in the Soret region.

## Abbreviations

Chl: chlorophyll
Chl_D1_/Chl_D2_: monomeric Chl on the D1 or D2 side, respectively
P_D1_ and P_D2_: individual Chl on the D1 or D2 side, respectively, which constitute a pair of Chl with partially overlapping aromatic rings (P_680_)
Phe_D1_ and Phe_D2_: pheophytin on the D1 or D2 side, respectively
PPBQ: phenyl *p*–benzoquinone
PSII: Photosystem II
Q_A_: primary quinone acceptor
Q_B_: secondary quinone acceptor
Tyr_D_: the tyrosine 160 of D2 acting as a side-path electron donor of PSII
Tyr_Z_: the tyrosine 161 of D1 acting as the electron donor to P_680_

## Acknowledgements

This work has been in part supported by (i) the French Infrastructure for Integrated Structural Biology (FRISBI) ANR-10-INBS-05, (ii) the Labex Dynamo (ANR-11-LABX-0011-01), (iii) the JSPS-KAKENHI Grant in Scientific Research on Innovative Areas JP17H064351 and a JSPS-KAKENHI Grant 21H02447 and (iv) the BBSRC grants BB/R001383/1, BB/V002015/1 and BB/R00921X.

## Notes

### Competing Interest Statement

The authors have declared no competing interest.

